# *In situ* evaluation of artificial selection and assisted gene flow for the advancement of flowering onset in *Lupinus angustifolius* L. (Fabaceae)

**DOI:** 10.1101/2025.07.23.666208

**Authors:** S. Sacristán-Bajo, A. García-Fernández, C. Lara-Romero, C. Poyatos, M.L. Rubio Teso, J. Morente-López, S. Prieto-Benítez, E. Torres, J.M. Iriondo

**Affiliations:** Grupo de Ecología Evolutiva (ECOEVO). Área de Biodiversidad y Conservación, Dpto. de Biología, Geología, Física y Química Inorgánica, ESCET, Universidad Rey Juan Carlos, C/ Tulipán s/n, 28933 Móstoles, Madrid, Spain; Instituto de Investigación en Cambio Global. Universidad Rey Juan Carlos, C/ Tulipán s/n, 28933 Móstoles, Madrid, Spain; Department of Plant Evolutionary Ecology, Institute for Ecology, Evolution and Diversity, Goethe-Universität, Max-von-Laue-Str, 60438, Frankfurt am Main; Grupo de Ecotoxicología y Contaminación del Aire, Departamento de Medio Ambiente, CIEMAT, Av. Complutense, 40, 28040 Madrid, Spain; Departamento de Biotecnología-Biología Vegetal, Universidad Politécnica de Madrid, Av. Puerta de Hierro, 2 – 4, 28040 Madrid, Spain

**Keywords:** Adaptive potential, biodiversity conservation, climate change, facilitated adaptation, natural conditions, restoration

## Abstract

Artificial selection and assisted gene flow represent promising conservation strategies for enhancing species’ adaptive capacity under accelerating climate change. We tested the application of artificial selection and assisted gene flow by evaluating progeny performance under field conditions, using *Lupinus angustifolius* L. (Fabaceae) as a model to advance flowering onset. Seeds were collected from four wild populations and two contrasting latitudes in Spain for parallel artificial selection and assisted gene flow three-year experiments. In the artificial selection treatment, we developed two early-flowering selection lines per population over three generations using both selfing and outcrossing. For assisted gene flow, we created F1 hybrids by crossing northern populations with southern pollen donors, then produced F2 and F3 generations through successive self-pollination. Finally, F3 seeds from this process were sown in autumn in a common garden under natural conditions, near the original population sites. Spring 2021 measurements of flowering onset and morphological traits revealed contrasting treatment effects. Artificial selection lines showed no significant phenotypic differentiation from controls across all measured traits, contradicting previous controlled-environment results. Conversely, assisted gene flow lines exhibited significantly earlier flowering and reduced shoot growth relative to controls, consistent with prior findings under controlled conditions. These results show greater efficacy of assisted gene flow over artificial selection for advancing flowering onset under natural conditions.

## Introduction

Climate change drives latitudinal and altitudinal range shifts in many organisms (Walther *et al*. 2002; Forero-Medina *et al*. 2011; Rubenstein *et al*. 2023), yet dispersal-limited plant species often cannot track suitable climate conditions (Engler *et al*. 2009). Alternatively, they may rely on *in-situ* adaptation to the novel environments through phenotypic plasticity or genetic adaptation (Jump and Peñuelas 2005; Chevin and Bridle 2025). However, accelerating climate change may exceed natural adaptive capacity, threatening population persistence (Urban 2015; Talukder *et al*. 2022). In this context, effective conservation requires assessing population adaptive capacity and identifying methods to enhance evolutionary potential.

Climate-threatened plant populations can enhance fitness through genetic variability that improves adaptation to new environmental conditions (Gomulkiewicz and Shaw 2013; Galland *et al*. 2025). Artificial selection and assisted gene flow represent two approaches for quantifying and increasing population adaptive potential (Hoffmann and Sgrò 2011; Aitken and Whitlock 2013; Anderson and Song 2020; Morente-López *et al*. 2021). Artificial selection preserves beneficial traits by selecting individuals with desired characteristics for breeding (Conner 2003, 2016), while assisted gene flow involves deliberately moving gametes, or individuals between populations to promote adaptation to current or future conditions (Aitken and Whitlock 2013; Whiteley *et al*. 2015; Morente-López *et al*. 2021). Both strategies constitute facilitated adaptation—human intervention in evolutionary processes through beneficial allele introduction or enhancement to help organisms adapt to new environmental pressures like those derived from climate change (Van Oppen *et al*. 2015; Humanes *et al*. 2021; Torres *et al*. 2023). These interventions can entail certain risks that need to be properly addressed (Torres *et al*. 2023). Artificial selection may reduce genetic diversity, potentially compromising evolutionary resilience (Whitt *et al*. 2002; Sheth and Angert 2016; Chen *et al*. 2017). Assisted gene flow can disrupt linkage disequilibrium by introducing maladapted alleles or break coadapted gene networks, leading to reduced fitness through maladaptation or outbreeding depression (Aitken and Whitlock 2013; Grummer *et al*. 2022). Additionally, both strategies may produce undesired correlated responses in other traits, constraining adaptation through evolutionary trade-offs (Sheth and Angert 2016; Sacristán-Bajo *et al*. 2023).

Experimental assessments of artificial selection and assisted gene flow have demonstrated their potential effects on population genetics and functional traits (Van Oppen *et al*. 2015; Morente-López *et al*. 2021; Poyatos *et al*. 2023). However, these studies were conducted under controlled conditions or as proof-of-concept experiments. Given the challenges of climate change, field testing in natural conditions is essential to evaluate whether these tools can effectively exert the desired phenotypic effects. Field experiments can replicate evolutionary processes difficult to observe in controlled environments (Garland Jr and Kelly 2006; Anderson and Wadgymar 2020). For instance, trade-offs that are absent or weakened under controlled conditions may represent major evolutionary constraints in natural settings. On the other hand, field experiments present interpretative challenges due to uncontrolled environmental factors that complicate causal inference (Garland and Rose 2009; Felton *et al*. 2024). Understanding these limitations is, therefore, crucial for predicting and interpreting evolutionary responses (Garland and Rose 2009).

Identifying climate-relevant adaptive traits amenable to intervention is essential for implementing these conservation strategies. Flowering onset critically determines plant reproductive success (Thomas *et al*. 2001; Forrest and Thomson 2010; Blümel *et al*. 2015) and exhibits strong genetic control (Franks *et al*. 2007; Franks and Hoffmann 2012). Climate change has already shifted flowering periods in temperate regions (Fitter and Fitter 2002; Büntgen *et al*. 2022), as organisms adjust their phenology to environmental changes (Bradshaw and Holzapfel 2008; Cohen *et al*. 2018). Consequently, flowering onset represents an ideal target trait for testing artificial selection and assisted gene flow strategies in plant climate adaptation.

This study aimed to test management tools to enhance species climate adaptation by evaluating artificial selection and assisted gene flow progeny under natural field conditions, targeting earlier flowering onset for potential conservation applications. This field experiment built on previous controlled *ex-situ* studies using *Lupinus angustifolius* L. (Fabaceae) populations from contrasting climatic zones (hereafter, northern and southern populations) where these treatments were implemented (Sacristán-Bajo *et al*. 2023). Under controlled conditions, artificial selection lines from northern populations showed significantly advanced flowering onset compared to controls, while southern populations showed no response. Selection lines maintained fitness but altered some vegetative traits (Sacristán-Bajo *et al*. 2023). Assisted gene flow lines (northern populations crossed with southern pollen) also advanced flowering onset and increased seed weight relative to controls, with additional vegetative effects (Sacristán-Bajo *et al*. 2025). The present field experiment addresses two objectives: validating that trait modifications persist under natural conditions and assessing whether these modifications enhance individual fitness.

Based on these controlled-condition results, we established a common garden experiment using treatment offspring under natural conditions near one of the northern populations. We hypothesized that artificial selection and assisted gene flow lines would maintain earlier flowering relative to controls under field conditions. In addition, since selective pressures are stronger under natural conditions, we also hypothesized that both artificial selection and assisted gene flow lines would experience changes in other fitness-related and vegetative traits compared to controls. Finally, since assisted gene flow produced greater flowering advancement than artificial selection under controlled conditions, we hypothesized this pattern would persist under natural conditions.

## Methods

### Study species and experimental design

*Lupinus angustifolius* L. is an annual legume that is extensively prevalent throughout the Mediterranean basin and has been introduced and grown as a crop in many other parts of the world (Castroviejo *et al*. 1996). In natural conditions, it inhabits well-drained acid or neutral soils, including disturbed environments such as roadsides or abandoned fields (Rhodes & Maxted, 2016). The species germinates in autumn and blooms between March and August depending on environmental conditions (Castroviejo *et al*. 1996). Its blue-purple flowers are zygomorphic and hermaphroditic, and mainly self-pollinate before the petals open (Wolko *et al*., 2011).

In 2016 we collected seeds from four populations spanning two climatically contrasting zones in Central and Southern Spain (Figure 1, Table S1). Seeds were harvested from a minimum of 98 maternal plants (hereafter, genotypes) per population, and subsequently sown and cultivated in a common garden environment at the CULTIVE facility (https://urjc-cultive.webnode.es/) at Rey Juan Carlos University (Móstoles, Madrid). An artificial selection experiment and a geneflow experiment were conducted in the common garden environment. For the artificial selection experiment, we established two breeding lines in 2017: an early flowering line (EFL) by selecting individuals from the first quartile of earliest flowering times, and a control line (CFL) by randomly selecting 25% of individuals. In spring 2018, we generated an outcrossed early flowering line (OUT) through manual crosses among EFL genotypes. The OUT line was subsequently self-crossed in spring 2019 to produce a segregating F2 generation (OUTS), which was then self-pollinated in 2020 to generate the F3 generation (OUTS2). Concurrently, both EFL and CFL lines were maintained through self-pollination, producing their respective F3 generations in 2020. For the gene flow experiment, we established an F1 gene flow line (GFL) in 2018 by manually pollinating plants from northern populations with pollen from southern populations, using the same control line (CFL) as in the artificial selection experiment. In 2019, the GFL was self-pollinated to generate an F2 segregating line (SPL), which was subsequently self-pollinated in 2020 to produce the F3 generation (SPL2). Complete methodological details, including growing conditions and hand-pollination protocols, are provided in Sacristán-Bajo et al. (2023, 2025), with the entire experimental procedure illustrated in Figure 2. The 2020 field experiment described in the present study utilized F3 seeds from four lines: CFL, EFL, OUTS2, and SPL2 from the northern populations: Zarapicos (PIC) and Zafrón (FRO).

**Figure 1.**
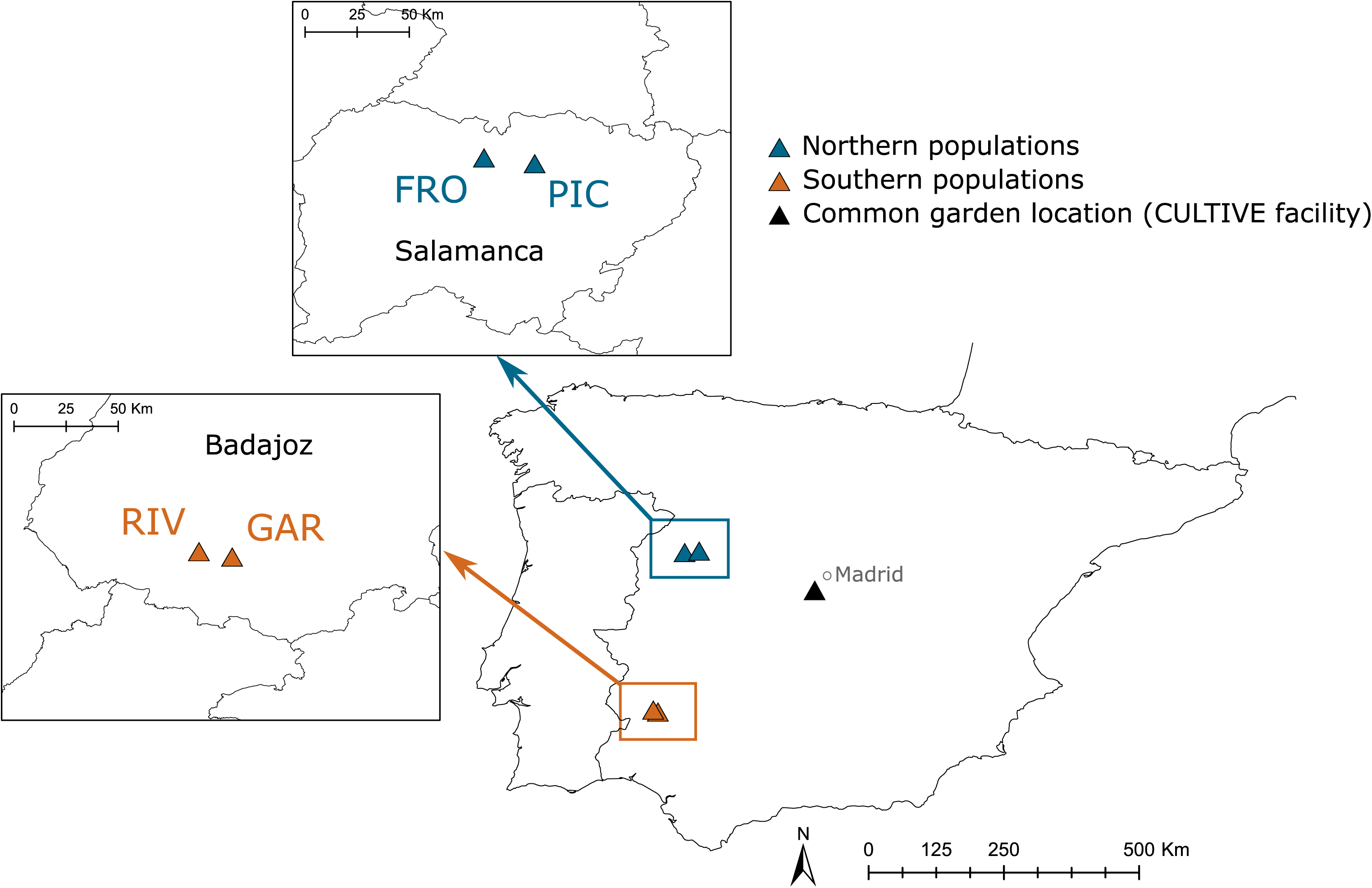
Location of the original populations of *Lupinus angustifolius* L., the site where the *ex-situ* common garden was implemented, and the plot used in the field experiment (common garden, natural conditions).

**Figure 2.**
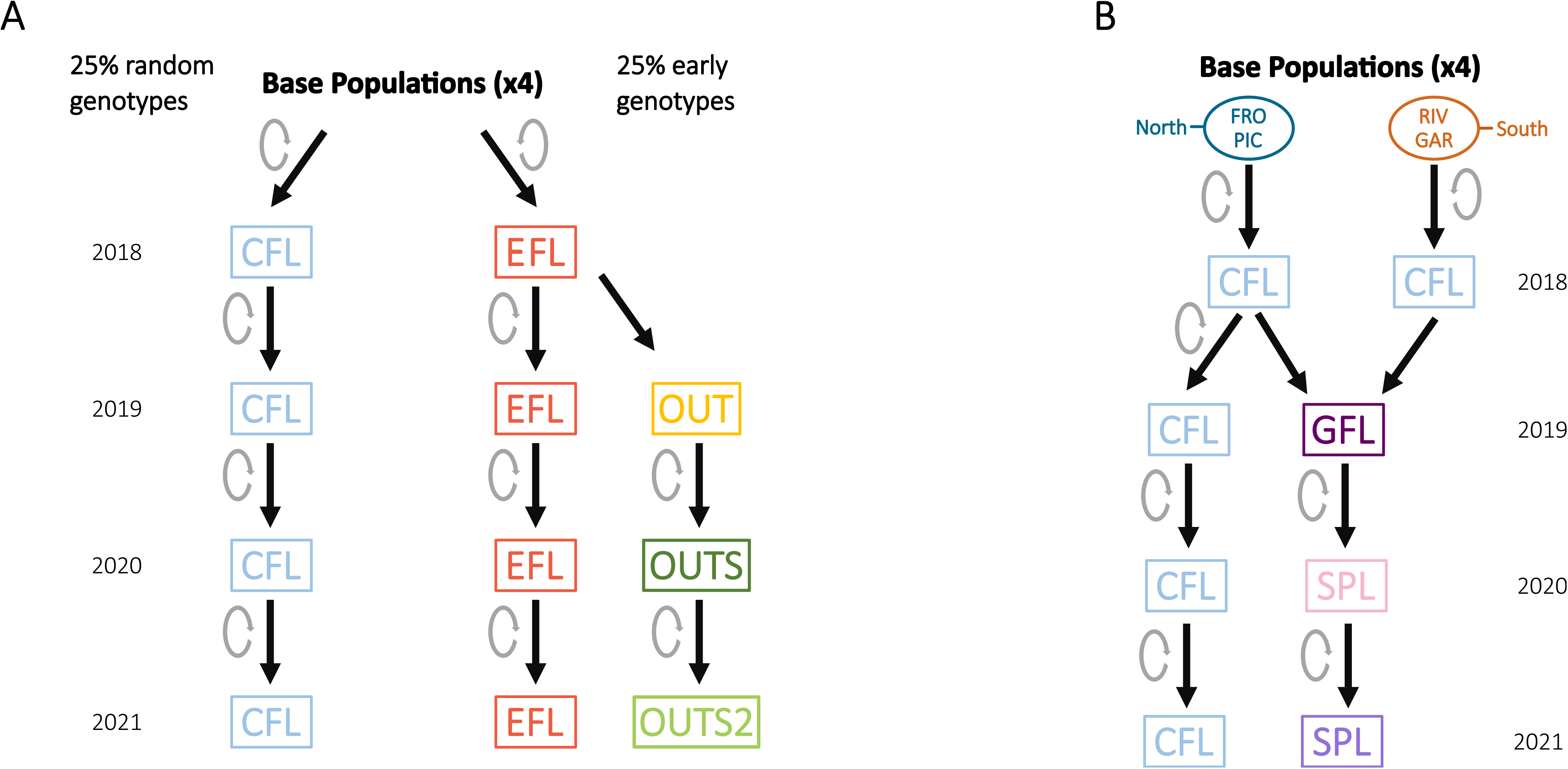
Breeding scheme showing line development across four *Lupinus angustifolius* populations. a) Artificial selection; b) assisted gene flow. Grey circles next to the arrows indicate self-crossing events. CFL: control flowering line; EFL: early flowering line (selfed); OUT: outbred line (crosses among EFL genotypes); OUTS: F2 generation from OUT self-pollination; OUTS2: F3 generation from OUTS self-pollination; GFL: F1 gene flow line (interpopulation crosses); SPL: F2 generation from GFL self-pollination; SPL2: F3 generation from SPL self-pollination.

In November 2020, we established the field experiment by sowing seeds from both northern population lines under natural conditions in cattle-grazed grassland at Zarapicos, Salamanca (Figures 1 and 3). The experimental design consisted of four randomized blocks, each containing seeds from both populations and all artificial selection and gene flow lines (Figure 3a). Within each block, we established 234 sowing points arranged in two parallel rows spaced 20 cm apart, with individual points separated by 20 cm in each row. Blocks were separated by one-meter corridors.

**Figure 3.**
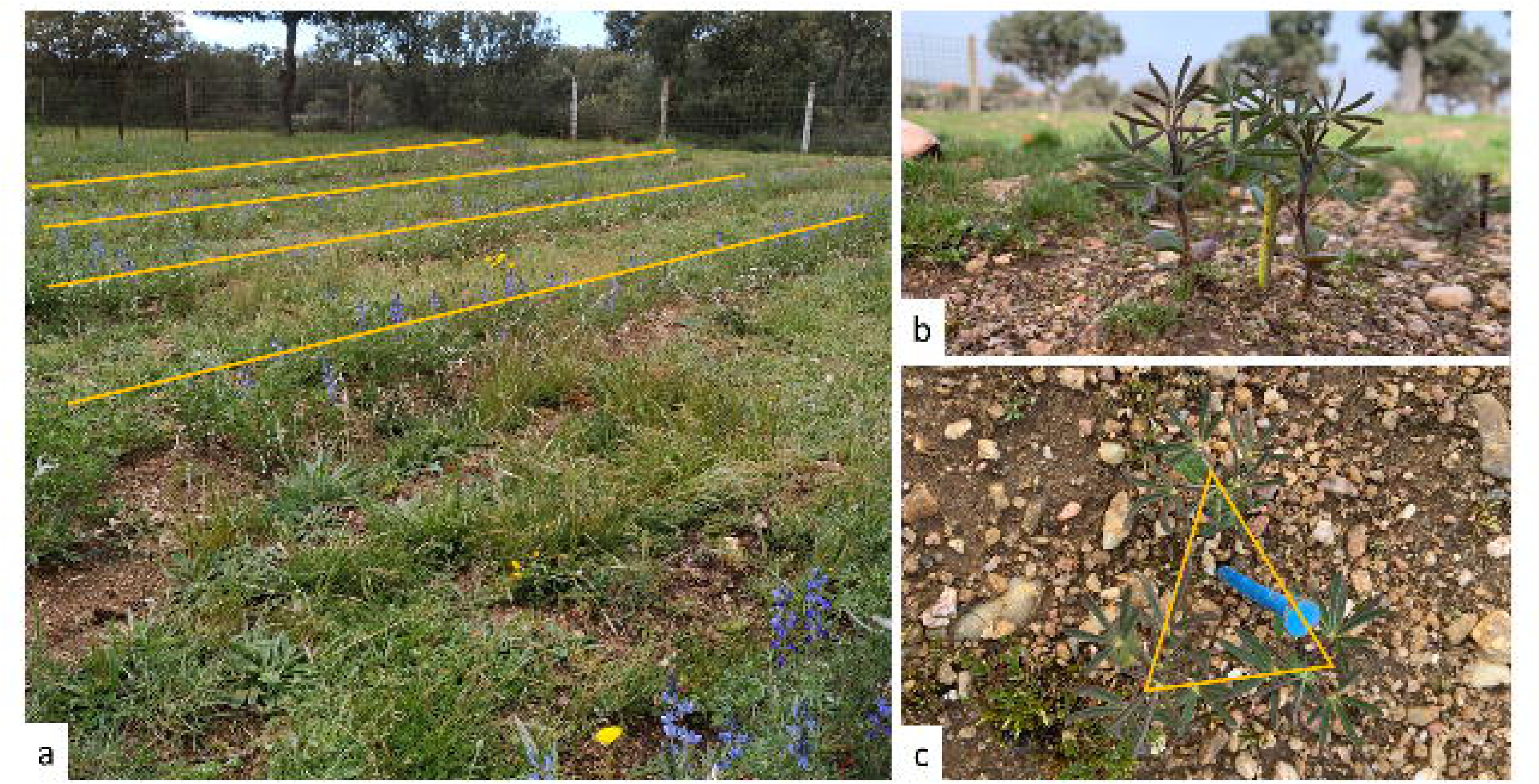
Experimental plot under natural conditions. a) In each block the sowing points of *Lupinus angustifolius* were arranged in two parallel lines that were 20 cm apart. b) Seedlings of *Lupinus angustifolius*. c) Triangular arrangement of three seedlings of a genotype at a sowing point.

At each sowing point, we manually prepared the soil with a hoe and sowed three pre-weighed, scarified seeds of the same genotype in a triangular arrangement (Figures 3b and 3c). The experiment included 2,808 total seeds: 36 seeds per genotype across 10 genotypes per treatment (8 genotypes for SPL2 in the FRO population) and four treatments per population. No supplemental irrigation or treatments were applied to maintain conditions representative of the natural populations which occur within a 20 km radius of the experimental site.

### Traits measurement

We monitored flowering onset, defined as the date of first flower opening, through tri-weekly observations throughout the flowering period. When plants flowered between observation dates, we inferred the precise flowering date based on flower developmental stage. The days to flowering onset were calculated as the number of days between sowing and flowering onset dates. Plant height was measured twice as the distance from ground level to the apex of the vegetative stem portion: initially on March 17, 2021 (flowering onset), and finally between May 14-18, 2021 (flowering conclusion). The difference between measurements provided shoot growth during the flowering period.

Following reproductive season completion in early June, when all flowers had developed into fruits, we harvested plants and quantified seed production per individual as a fitness proxy. We also measured total plant dry biomass at harvest.

### Statistical analyses

We conducted separate analyses comparing artificial selection and assisted gene flow treatments against the same control treatment. All analyses were performed in R statistical environment version 4.1.1 (R Core Team 2021) using flowering onset, shoot growth, biomass, and seed number as response variables. We fitted generalized linear mixed models (GLMMs) for flowering onset and linear mixed models (LMMs) for all other traits using the glmmTMB package version 1.1.3 (Brooks *et al*. 2017). Models assumed Poisson and Gaussian distributions for flowering onset and remaining traits, respectively. All models included *line* and *population* as fixed effects, *genotype* (maternal plant) as a random effect, and *germinated seed weight* as a covariate. Model assumptions were verified through diagnostic residual plots.

The significance of each fixed effect was evaluated using the *Anova* function from the R package version 3.0-11 (Fox and Weisberg 2018). We calculated R² values using the *r*.*squaredGLMM* function from the package *MuMIn* version 1.47.1 (Nakagawa and Schielzeth 2013). Posterior mean values, standard errors and 95 % confidence intervals for the different traits and lines were calculated using the *emmeans* package version 1.6.3 (Lenth 2023). Correlations between the flowering onset and the other variables within the control line were plotted using the *corrplot* function from the package *corrplot* version 0.90 (Wei *et al*. 2017).

## Results

### Artificial selection experiment

Flowering onset did not differ significantly among artificial selection lines (EFL, OUTS2) and the control (CFL) across populations (χ² = 3.512, p = 0.173, df = 2) (Figure 4a; Tables S2, S3). Similarly, no significant differences were detected between artificial selection and control lines for seed number (χ² = 5.35, p = 0.07, df = 2), biomass (χ² = 0.49, p = 0.78, df = 2), or shoot growth (χ² = 3.98, p = 0.14, df = 2) (Figures 4b–d; Tables S2, S3).

**Figure 4.**
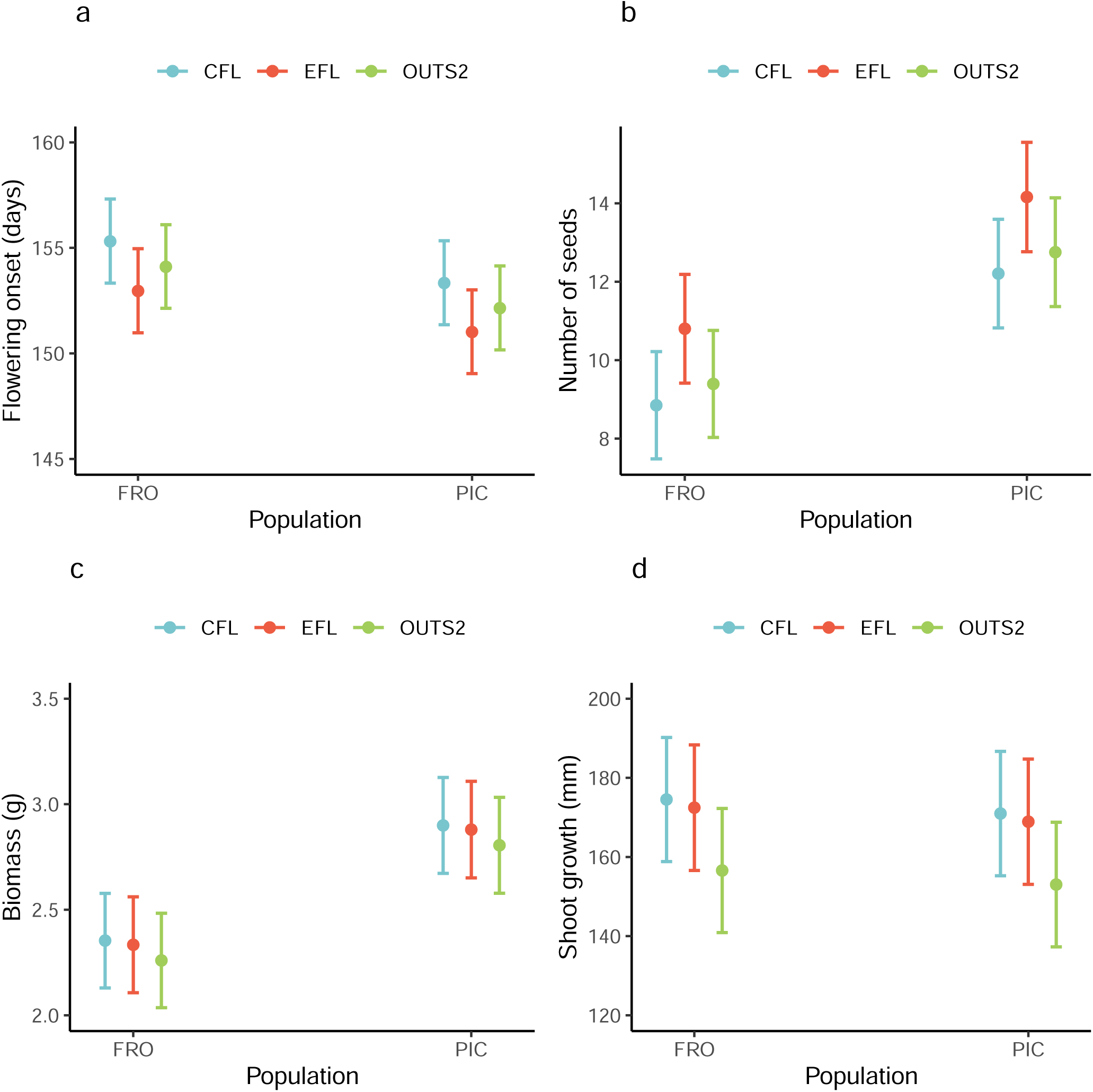
Effects of artificial selection lines (EFL and OUTS2) on *L. angustifolius* traits. a) flowering onset, b) number of seeds, c) biomass, d) shoot growth. CFL: control flowering line; EFL: early flowering line (selfed); OUTS2: F3 generation from two generations of OUT self-pollination.

In contrast, the FRO and PIC populations differed significantly in seed number (χ² = 22.87, p < 0.001, df = 1) and biomass (χ² = 22.48, p < 0.001, df = 1), with PIC plants producing more seeds and greater biomass than FRO plants (Figures 4b, 4c; Table S4). Posterior means, standard errors, and 95% confidence intervals are provided in Table S5.

### Gene flow experiment

The SPL2 line flowered significantly earlier than the control (CFL), with a 10-day advance in the FRO population and 4-day advance in the PIC population (χ² = 17.12, p < 0.001, df = 1) (Figure 5a; Tables S4, S6, S7). SPL2 individuals also exhibited reduced shoot growth relative to controls, averaging 25 mm less in FRO and 21 mm less in PIC populations (χ² = 8.26, p = 0.004, df = 1) (Figure 5b; Tables S4, S6, S7). Fixed effects explained 10.2% and 11.6% of variation in flowering onset and shoot growth, respectively, while random effects explained 5% and 39.6% (Table S6).

**Figure 5.**
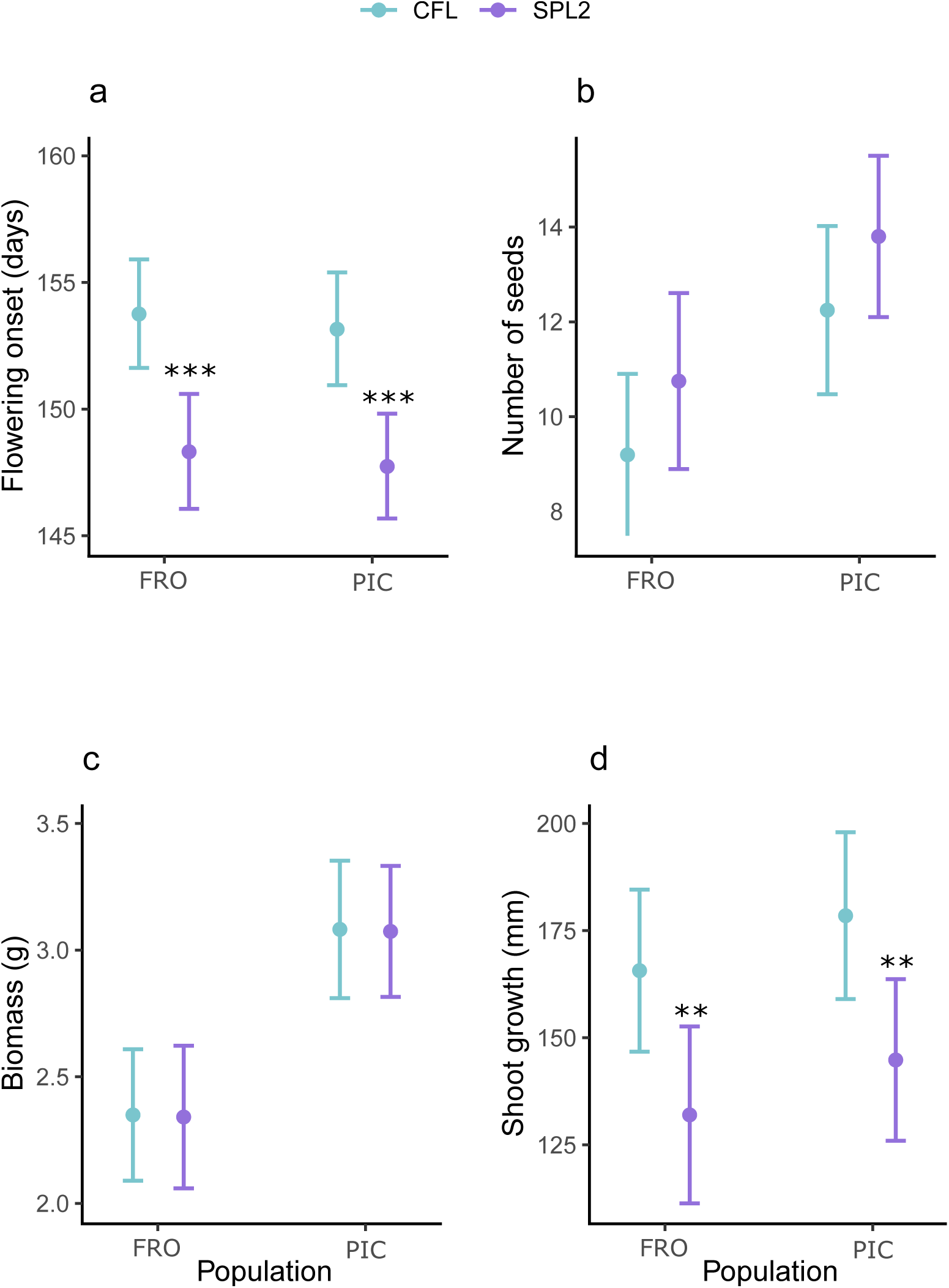
Effect of the assisted gene flow line (SPL2) on *L. angustifolius* traits. a) flowering onset b) number of seeds, c) biomass, d) shoot growth. CFL: control flowering line; SPL2: F3 generation from two generations of gene flow line (GFL) self-pollination.

No significant differences between lines were detected for seed number (χ² = 2.16, p = 0.14, df = 1) or biomass (χ² = 0.003, p = 0.96, df = 1). However, both traits differed significantly between populations (seed number: χ² = 8.75, p = 0.003, df = 1; biomass: χ² = 21.79, p < 0.001, df = 1) (Figures 5c, 5d; Tables S6, S7). Consistent with the artificial selection experiment, PIC plants produced more seeds and greater biomass than FRO plants (Figures 5c, 5d; Table S4). Posterior means, standard errors, and 95% confidence intervals are provided in Table S8.

### Correlations between traits

Flowering onset correlated significantly with seed number, biomass, and shoot growth (Figure S1). In both populations, earlier flowering was associated with increased seed production (r_FRO_ = -0.49, r_PIC_ = -0.36) and higher biomass (r_FRO_ = -0.34, r_PIC_ = -0.50) (Figure S1a,b). Conversely, shoot growth showed opposite sign correlations with flowering onset between populations: negative in FRO (r = -0.15) but positive in PIC (r = 0.18) (Figure S1a,b).

## Discussion

Artificial selection and assisted gene flow differed markedly in their effectiveness under natural conditions. Artificial selection lines showed no significant changes in flowering onset or other measured traits relative to controls. In contrast, the assisted gene flow line advanced flowering significantly in both populations, albeit with reduced shoot growth.

Notably, assisted gene flow results were consistent between natural and controlled conditions (Sacristán-Bajo *et al*. 2025), whereas artificial selection effects observed under controlled conditions (Sacristán-Bajo *et al*. 2023) were not found in the field. These contrasting patterns suggest genotype-by-environment interactions influence artificial selection responses and indicate that different mechanisms may govern flowering time expression under each approach.

Our findings emphasize the critical importance of field validation across diverse environmental conditions when evaluating facilitated adaptation strategies for conservation and restoration applications (Anderson and Wadgymar 2020; Sammarco *et al*. 2022).

### Effects of artificial selection

Neither artificial selection line (EFL, OUTS2) significantly advanced flowering onset compared to controls, despite drier-than-normal field conditions (Table S1). All lines flowered at 154-155 days after sowing, contrasting sharply with controlled-condition results where selection lines flowered 115-136 days after sowing and significantly earlier than controls in northern populations (Sacristán-Bajo *et al*. 2023). The delayed flowering under field conditions might be explained by cooler field temperatures relative to controlled environments and the existence of accumulated heat requirements for flowering initiation (Galán *et al*. 2001). However, this thermal explanation cannot account for the loss of line differences observed under controlled conditions, particularly given uniform mortality and germination rates across lines and populations. The differences found in the results between both experiments point to a strong genotype-by-environment (G x E) interaction, where phenotypic plasticity varies among genotypes due to differential gene expression across environments (Clausen *et al*. 1940; Basford and Cooper 1998; Wei *et al*. 2025).

Such G x E interactions have critical implications for predicting selection responses, as environmental changes can alter expected outcomes (Matesanz and Ramírez-Valiente 2019). In this sense, the greater selection differentials observed in northern versus southern populations (Sacristán-Bajo *et al*. 2023) suggest a more reduced selective pressure for early flowering in northern populations under natural conditions. Additionally, it should be noted that the OUTS2 line sown in the field experiment was genetically different from the OUT and OUTS lines sown under controlled conditions. The hybrids and their descendants (OUT and OUTS respectively) tested under controlled conditions had higher heterozygosity than the field-tested OUTS2 line, because the latter experienced two generations of self-pollination. This could also potentially contribute to the differential responses observed.

Additional factors not considered may influence these results, including vernalization requirements (Adhikari *et al*. 2012; Jiménez – López *et al*. 2024), substrate composition (Ausin *et al*. 2005), and biotic interactions under varying conditions (Ge *et al*. 2025). Vernalization promotes flowering in many species through exposure to low temperatures (Adhikari *et al*. 2012), and while lupin flowering is typically controlled by vernalization, some *Lupinus angustifolius* cultivars have been selected for vernalization independence (Gladstones and Hill 1969; Rahman and Gladstones 1974), evidencing the existence of vernalization-independent genotypes in wild populations.

Under controlled conditions with limited cold exposure, artificial selection may have favored vernalization-independent genotypes that flower early without cold treatment, while control lines likely retained both vernalization-dependent and -independent genotypes. In the field, where plants experienced natural cold periods, both genotype classes would synchronize flowering onset, eliminating observable differences between selection and control lines. This vernalization hypothesis could also explain the contrasting results between controlled and field experiments.

### Effects of assisted gene flow

Unlike artificial selection, assisted gene flow effects observed under controlled conditions persisted under natural conditions. Remarkably, differences between lines remained detectable after two generations of selfing following interpopulation crosses, despite more reduced heterozygosity. This consistency reinforces the strong genetic basis of assisted gene flow modifications, likely involving novel allele incorporation (Sacristán-Bajo *et al*. 2025) and demonstrates the potential for providing genetic variation necessary for climate adaptation (Aitken and Whitlock 2013).

Recent simulations suggest assisted gene flow may cause short-term harm (10-20 generations) through outbreeding depression, with polygenic traits requiring longer timeframes for beneficial effects (Grummer *et al*. 2022). However, our results show no detrimental effects in early generations, despite flowering time’s polygenic nature.

As expected, flowering onset modification induced correlated trait changes. SPL2 plants exhibited reduced shoot growth but maintained reproductive success and biomass, consistent with controlled-condition results. The growth reduction likely reflects resource allocation trade-offs (Reich 2014; Sobral 2021). While controlled experiments revealed increased seed weight without changes in seed number (Poyatos *et al*. 2023), seed weight could not be measured under field conditions.

Notably, correlation analyses in control lines showed that earlier flowering correlated with increased seed production and biomass, contrasting with the lack of changes in reproductive success in SPL2. This discrepancy may reflect environment-dependent genetic correlations, where favorable trait combinations in certain environments may not be advantageous in others (Sheth and Angert 2016; Sobral 2021; Sammarco *et al*. 2022).

### Artificial selection vs. assisted gene flow

Assisted gene flow proved more effective than artificial selection for advancing *L. angustifolius* flowering under natural conditions. While artificial selection achieved greater flowering advances than assisted gene flow under controlled conditions in northern populations (Sacristán-Bajo *et al*. 2023, 2025), these effects disappeared in the field, whereas assisted gene flow effects persisted (Sexton *et al*. 2024).

These contrasting patterns suggest different underlying mechanisms. Assisted gene flow likely modifies flowering through direct genetic changes, introducing early-flowering alleles from southern into northern populations, as confirmed by genomic analyses (Sacristán-Bajo *et al*. 2025). Conversely, artificial selection effects may operate through regulatory processes—epigenetic mechanisms or regulatory gene changes—that are more environmentally sensitive (Mylne *et al*. 2004; Ausin *et al*. 2005; Sammarco *et al*. 2022).

The SPL2 gene flow line contains in average 50% southern and 50% northern genomic content, which advances flowering but reduces northern genetic identity and potentially compromises adaptation to unmeasured environmental factors. While natural selection may eliminate these maladaptive effects over time (Aitken and Whitlock 2013; Grummer *et al*. 2022), backcrossing combined with selection strategies could accelerate recovery of local genetic identity while preserving desired traits.

In conclusion, flowering onset is a complex trait involving the interaction of multiple genes and diverse genetic and epigenetic mechanisms (Blümel *et al*. 2015; Chen *et al*. 2018), precluding simple interpretation or prediction of responses to different interventions.

### Final remarks

Our experiments demonstrate that assisted gene flow effects are both stronger and more environmentally stable than artificial selection responses. These findings underscore the critical importance of multi-environment common garden experiments for evaluating facilitated adaptation strategies. Since our field experiment was conducted under cooler conditions than controlled environments, additional field trials at warmer temperatures would complement the present results to simulate projected climate warming scenarios.

Climate change poses unprecedented challenges for biodiversity conservation and ecological restoration (Hancock *et al*. 2013). Studies of this kind are essential for assessing population adaptive potential and identifying optimal strategies to enhance species survival under changing environmental conditions.

## Acknowledgements

We thank all field work participants, including AdAptA-lab members and undergraduate students. We especially thank Carlos Ingala for all his efforts and support. This research was supported by the EVA project (CGL2016-77377-R) and DACWIRE project (PID2021-127841OA-I00 funded by 399 MICIU/AEI /10.13039/501100011033/ and by ERDF A Way for Europe) from the Spanish Ministry of Science and Innovation.

## Data availability statement

Data associated with this study are made available in the figshare data repository: 10.6084/m9.figshare.29606180

